# Dual nuclease single-cell lineage tracing by Cas9 and Cas12a

**DOI:** 10.1101/2024.05.27.596125

**Authors:** Cheng Chen, Yuanxin Liao, Miao Zhu, Li Wang, Xinran Yu, Meishi Li, Guangdun Peng

**Affiliations:** Center for Cell Lineage and Development, CAS Key Laboratory of Regenerative Biology, Guangdong Provincial Key Laboratory of Stem Cell and Regenerative Medicine, GIBH-HKU Guangdong-Hong Kong Stem Cell and Regenerative Medicine Research Centre, China-New Zealand Joint Laboratory on Biomedicine and Health, Guangzhou Institutes of Biomedicine and Health, Chinese Academy of Sciences, Guangzhou 510530, China; University of Chinese Academy of Sciences

**Author notes:** These authors contributed equally (Cheng Chen, Yuanxin Liao, Miao Zhu).

**Keywords:** single cell lineage tracing, Cas9, Cas12a, inter-site deletion, entropy

## Abstract

Single-cell lineage tracing based on CRISPR/Cas9 gene editing enables the simultaneous linkage of cell states and lineage history at a high resolution. Despite its immense potential in resolving the cell fate determination and genealogy within an organism, existing implementations of this technology suffer from limitations in recording capabilities and considerable barcode dropout. Here, we introduce DuTracer, a versatile tool, which utilizes two orthogonal gene editing systems to record deep cell lineage history at single-cell resolution in an inducible manner. DuTracer shows the ability of enhanced lineage recording with minimized target dropouts and deepened tree depth. Application of DuTracer in mouse embryoid bodies and neuromesodermal organoids illustrates the lineage relationship of different cell types and proposes potential lineage-biased molecular drivers. Moreover, we have developed an entropy-based approach to quantify the lineage recording ability of DuTracer in cell differentiation models. Together, DuTracer facilitates the precise and regulatory interrogation of cellular lineages of complex biological processes.

## INTRODUCTION

Charting cell states and dissecting cell lineage are crucial for understanding the mechanisms that govern cell fate determination, disease progression, as well as tissue injury and repair ^1–3^. The rapid advancements in single-cell sequencing technology have revolutionized our ability to precisely document the RNA molecules in individual cells ^4–7^, providing a vivid profiling of cell status. Trajectory inference analysis based on single-cell sequencing data further facilitates the characterization of cell state transitions ^8^. However, inferred cell trajectories may not represent the authentic lineage history of cells, as assumptions based on transcriptome similarity or RNA dynamics may not hold in real biological contexts ^9–12^. As the gold standard for determining cell lineage, genetic tracing can provide accurate lineage histories between cells but is restricted by limited read-out ^13–16^. By combining single-cell RNA sequencing with barcode-based lineage tracing, cutting-edge single-cell lineage tracing technology can simultaneously capture both the cell states and lineage histories, thus holding immense promise to unravel developmental and disease mechanisms ^17–23^.

Evolvable cellular barcodes inherently record the continuous lineage relationships of longer temporal range and higher complexity. Numerous studies have explored various strategies to generate barcodes via CRISPR editing, with the primary objective of achieving high information content ^20,24,25^. To adapt to popular high-throughput scRNA-seq methods, most approaches prefer concatenating multiple CRISPR/Cas9 targets into a tandem array of less than 250 bp ^26–29^. However, this arrangement of targets often fails to fully utilize the recording ability of each element, as the compact multiple targets are prone to deleterious inter-site deletions spanning multiple sites upon the simultaneous cutting of Cas9 protein. This greatly reduces the usable targets and decreases the resolution of the lineage tree. For instance, by arranging ten targets together, state-of-the-art CARLIN and its successor DARLIN tracing methods can generate millions of barcodes and record the clonal origin of hematopoietic stem cells ^25,28^. However, the inter-site deletion and exhaust of targets still hinder the tree-level construction of lineage within individual clones. Another consequence of large inter-site deletions is that they may even remove previously generated in-between insertions or deletions (indels), leading to potential severe errors in reconstructing the phylogenetic relationship between cells ^25,27^.

To circumvent the issue of large inter-site deletions, several methods have been developed. A common implementation is to reduce the number of targets per array; however, this may diminish the total barcode diversity per array and require increasing the copies of arrays ^3,30,31^. An alternative strategy is to decrease the chance of simultaneous cutting, either achieved by attenuating the cutting activity of a single nuclease, separating the editing timing of different targets by Cas9 induction ^30^, or introducing self-evolving elements scarred by a homing editing system ^32,33^. Nevertheless, the current practice of these methods suffers from either extended editing intervals or limited flexibility to sufficiently record a particular developmental process.

Barcode diversity is an important metric for evaluating barcode strategies in single-cell lineage tracing ^24^. As the barcode diversity of CRISPR indels is shaped by the interplay of cell division and gene editing, this metric only reflects the number of uniquely labeled clones captured by a designated sequencing. In addition to distinguishing cells, evolvable CRISPR barcodes are expected to record major cell fate determination or cell state transition events during a cell differentiation process ^21^, which is not easily revealed by barcode diversity alone. While many methods exist for visualizing and calculating lineage coupling of different cell types, there is a lack of proposed metrics to quantitatively evaluate the extent of cell differentiation events embedded in various barcoding strategies.

Here we present DuTracer, a single-cell lineage tracing technology leveraging two orthogonal CRISPR gene editing systems, Cas9 and Cas12a, to record cellular phylogenies. We demonstrated that DuTracer minimized inter-site deletions in HEK293T cell lines, mouse embryonic stem cells, mouse embryoid bodies, and mouse neuromesodermal organoids. By separating the expression of these two gene-editing nucleases, DuTracer enabled the building of a deeper lineage tree when compared with a single nuclease tracer at an equivalent number of targets. Using DuTracer as a system for evaluating gene editing properties within the same cell, we discovered that a defined barcoding system might inherently have constrained barcoding capacity, even when multiple insertion copies of targets are present. We established an information-theory-based method to summarize the recorded cell fate determination events for different cell types and quantify the extent of cell differentiation distinguished by barcoding processes. We further showed that the DuTracer system could help to reveal the heritable molecular features that marked the differentiation of second heart fields in a mouse embryonic body model. Finally, we demonstrated that DuTracer could facilitate the identification of candidate molecular drivers that regulate cell fate choices of neuromesodermal progenitors. Collectively, DuTracer offers a versatile tool for unraveling the complexities of cell lineage and fate determination to facilitate the understanding of developmental biology and diseases.

## RESULTS

### DuTracer displays minimal inter-site deletions

Barcode deletion causes information loss and is problematic in reconstructing phylogenetic trees. We specifically focus on inter-site deletions spanning more than 2 targets (defined as harmful inter-site deletions), as they may lead to severe errors by removing in-between records (Figure S1A, B). By evaluating popular Cas9-based barcoding strategies that utilize an array of multiple targets, we found harmful inter-site deletions accounted for 20-40% of the total generated alleles (Figure S1C, D). Theoretically, having fewer targets concatenated within the same genomic locus would reduce the frequency of harmful inter-site cutting. However, we still observed frequent inter-site deletions even in a three-target lineage recording system from Chan et al ^30^ (Figure S1E-H). To mitigate the occurrence of such information-loss inter-site deletions, we explored further by reducing the number of tandem targets to only two for a single nuclease. In the meanwhile, we introduced orthogonal sets of targets from two different nucleases to attain high barcode diversity. As a result, we devised a lineage-tracing strategy named DuTracer (Figure 1A).

**Figure 1.**
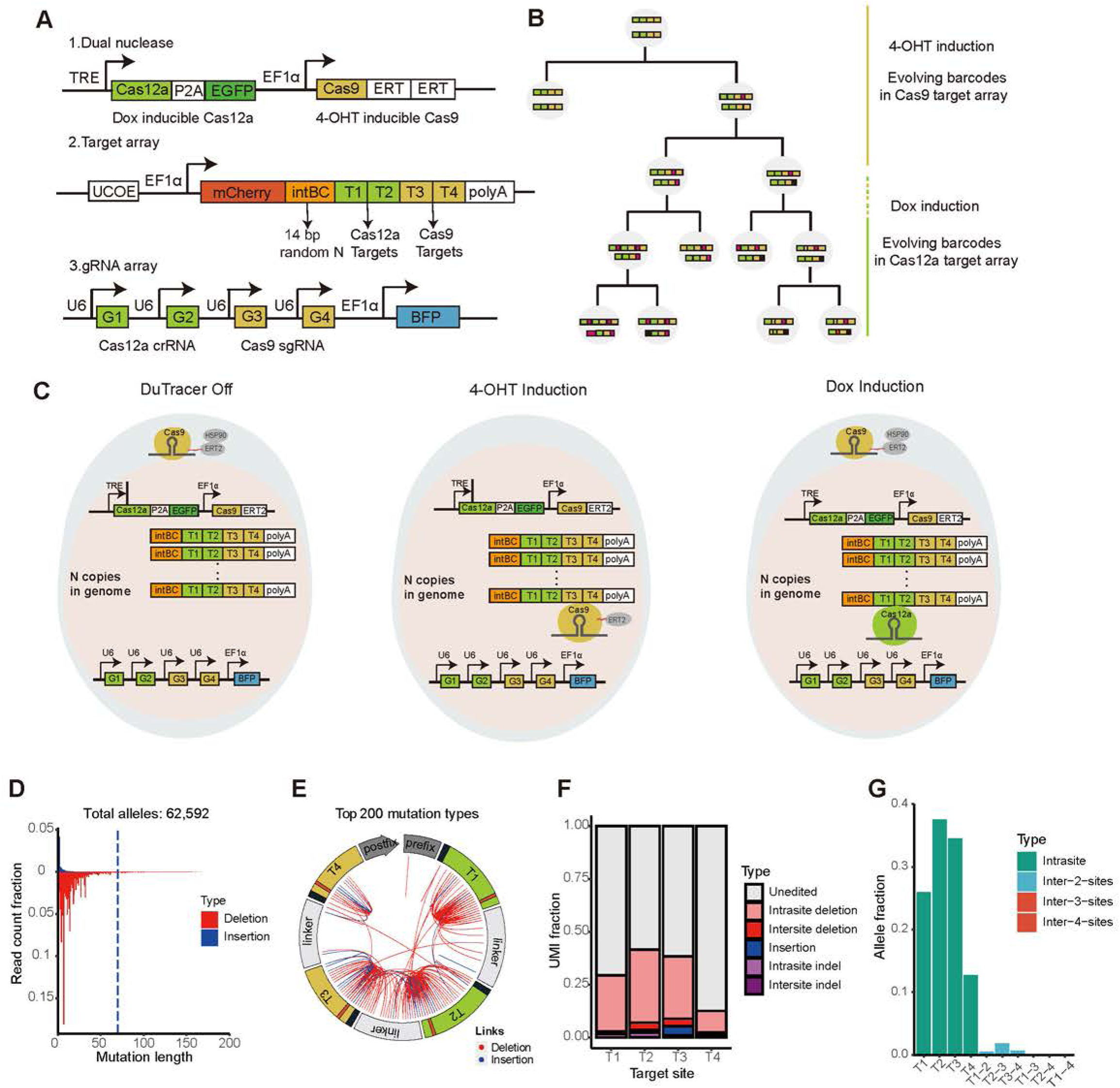
DuTracer generates minimal inter-site deletions. (A) Schematic of DuTracer system. Inducible Cas12a protein and inducible Cas9 components are placed within a PiggyBac backbone. The DuTracer target array sits in the 3’ UTR of mCherry and comprises two Cas12a sites and two Cas9 sites that perfectly match their corresponding gRNAs or crRNAs. Human U6 promoters are used to drive the expression of the two Cas12a crRNA and two Cas9 gRNAs. (B) Illustration of the lineage tree reconstruction through separate gene editing events. (C) Separate induction of nuclease activity using 4-OHT and doxycycline (Dox). (D-G) Editing patterns of DuTracer in eight mouse bulk ESC samples. (D) The length distribution of indels. Deletions with lengths greater than 70 bp (indicated using a dotted line) are considered as information-loss inter-site deletions. (E) Circos plot of top 200 indels. Indicated with circular rectangles, target sites and their surrounding sequence are counter-clock-wise arranged. The direct cut site within a target site is indicated by a small red rectangle, and the PAM region is illustrated with a black rectangle. The length of a circular rectangle is proportional to the number of base pairs of elements. Red lines display the starting and ending bp of deletion. Each blue line indicates the starting point of insertion and its insertion length, which is the offset between the endpoint and the starting point. (F) The read count fraction of different mutation types. (G) The allele fraction of different intrasite and inter-site indels. Intrasite, mutations within a single target. Inter-2-site and inter-3-site, deletions spanning two adjacent targets or three targets, respectively. Potential information-loss inter-site deletions are indicated with red color.

DuTracer employs two orthogonal nucleases, controlled by two different inducible systems (Tet-On and 4-OHT-ERT) ^34^ to separate the editing events of Cas9 and Cas12a (Figure 1 B, C). Additionally, to increase the total number of CRISPR targets per cell, we inserted multiple copies of target arrays into a cell. A 14 bp random DNA sequence is placed before the targets, serving as integration barcodes (intBCs) to distinguish different integration sites of various copies and reveal clonal relationships among cells ^31^ (Figure 1A). The non-overlapped cutting of Cas9 and Cas12a would minimize the target dropout and allow for a more controllable recording of lineage history.

To analyze the mutation pattern of DuTracer, we tested this system in HEK293T cells and mouse embryonic stem cells (ESCs) using different recording strategies. Specifically, for HEK293T cells, we initiated the Cas12a-directed tracking followed by Cas9 expression. In contrast, for mouse ESCs, Cas9-based recording preceded Cas12a activation. Albeit the difference in indel frequencies, both HEK293T cells and mouse ESCs exhibited thousands of mutated alleles, and indels predominantly occurred within individual target sites. We observed minimal inter-site deletions that span more than two target sites, representing a substantial improvement compared to the three-target design (Figure 1D-G and Figure S1I-L). Collectively, our experiments demonstrate that DuTracer greatly reduces the occurrence of information-loss inter-site deletions commonly encountered in tandem target arrays.

### Evaluation of DuTracer lineage reconstruction ability in HEK293T

To evaluate the lineage reconstruction ability of DuTracer in a simple cell division model, we generated a tracing HEK293T cell line with a high number of recording sites (Figure S2A). Starting with a population of 10 cells, we induced Cas12a expression for 22 days, followed by an additional 6 days of Cas9 expression (Figure 2A). We enriched the target cells by fluorescence-activated cell sorting and conducted scRNA-seq and amplicon-seq of target regions to reconstruct the cell lineages. We found the captured cells were inserted with an average of 22 intBCs (88 targets) and dominated by a large clone containing nearly 2,800 cells (Figure 2C and Figure S2B). One potential reason for the different sizes of the clones may be the differential level of barcode silencing (Figure S2C). DuTracer exhibited a multitude of mutation alleles, and a detailed assessment of the indels demonstrated that most indels occurred within individual targets, corroborating previous findings (Figure S1D-G). To further explore the lineage relationships, we selected the dominant clone and constructed a complex phylogenetic tree with an average depth of five layers (Figure 2B).

**Figure 2.**
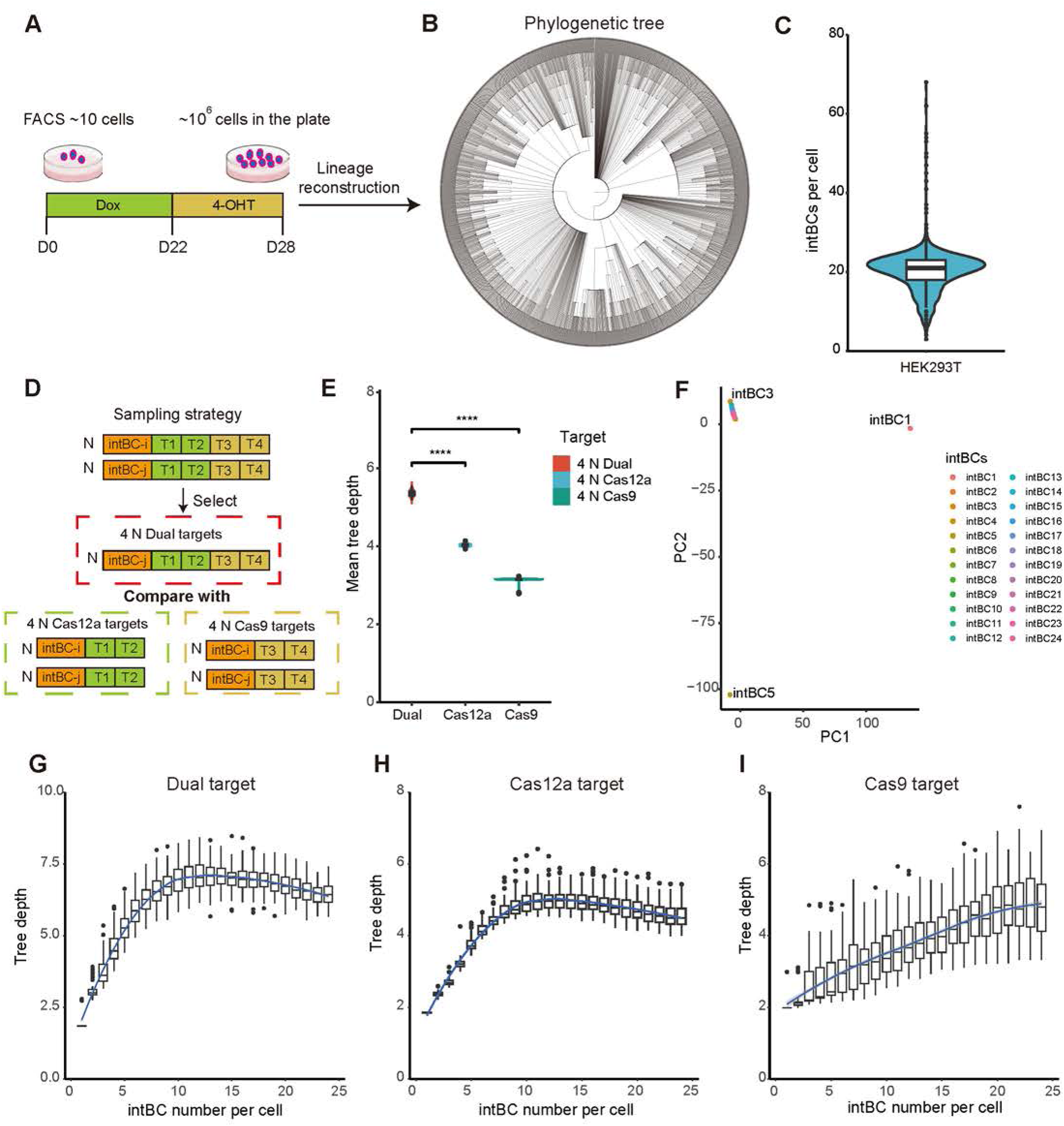
Evaluation of DuTracer lineage reconstruction ability in HEK293T. (A) Illustration of the experimental design. (B) The phylogenetic tree reconstructed from the largest clone. (C) Violin plot of the intBCs per cell. Cells from all clones were considered. (D) Illustration of the sampling strategy to obtain the same number of targets. N, the maximal even copy number of intBC pairs within a cell. For simplification, only a pair of target arrays is shown. (E) Simulated average tree depths under the same number of targets for dual targets, Cas9 targets, and Cas12a targets (unpaired two-tail t-test is indicated: ****p<0.0001). Each selection strategy was simulated 100 times. (F) Principal component analysis of the indel patterns of 24 intBCs within the largest clone. The UMI frequency of each intBC is used to order the intBCs. For example, intBC1 has the highest UMI counts, whereas intBC24 has the lowest UMI counts. (G-I) The relationship between average tree depth and the copy number of target arrays for dual targets (G), Cas12a targets (H), and Cas9 targets (I). Each copy number was sampled 100 times and variance is revealed by the boxplot.

A complete phylogenetic tree reflects the cell division events recorded by all accumulated mutations from both nucleases, but can also be tested parallelly against independent recorders. In our HEK293T experiment, we detected all the indels generated within four targets. We could construct a phylogenetic tree by combining indel information from all targets. Alternatively, we could create a tree using Cas9 targets information alone, and vice versa for the Cas12a tree. The dual tracing system allows us to evaluate the tree structure separately *in silico*. Through analyzing the given mutations in simulated target combinations (Figure 2D; Methods), we demonstrated that dual nuclease gene editing captured a deeper lineage hierarchy compared to single nuclease gene editing when using the same total number of targets (Figure 2E, Figure S2E-G). As expected, separate editing by two nucleases enables mutation events to occur across more cell division events.

Our DuTracer features two gene editing systems, providing a unique opportunity to interrogate critical lineage tracing challenges within the same cell by leveraging complementary barcoding processes. The copy number of target arrays is an important parameter for lineage reconstruction, as it determines the total number of targets available for gene editing. As intBC is located next to the targets, it serves as a landmark for anchoring targets. Our analysis revealed that targets in most intBC loci exhibited similar mutation profiles, while targets in a few intBC loci harbored distinct mutation patterns (Figure 2F). For example, mutations that occurred in Cas9 targets within the intBC1 locus were rare, whereas Cas9 targets within the intBC2 and intBC3 loci showed similar mutation patterns (Figure S2 H-J). To examine the impact of target array copy number on lineage recording depth, we conducted simulation experiments that randomly sampled targets containing 1 to 24 intBC loci per cell. The results demonstrated that the depth of the reconstructed tree reached a plateau at 10 intBCs (Figure 2G). Analyzing the saturated curve of Cas9 and Cas12a targets separately, we found that the rapid gene editing by Cas12a reached its plateau at 10 intBCs, while slower gene editing Cas9 did so at 22 intBCs (Figure 2H-I). This suggests that targets with different mutation rates may require varying copy numbers of target arrays to fully exploit their recording capacity. Importantly, our results indicate that while increasing the copies of tandem target arrays can increase the total number of recording sites per cell, there may be an upper-bound recording capacity. Beyond a certain level of copy numbers, an excess of targets may lead to redundant indels, failing to distinguish between cells.

### DuTracer has constrained barcoding capacity in mouse embryoid bodies

Next, we sought to test the lineage recording ability of DuTracer for a differentiation model in which mouse embryoid bodies (EB) produce diverse cell types within two weeks (Figure S3A)^35^. To delineate the barcoding capacity of each nuclease and reveal the consecutive lineage segregation during a complex biological cascade, we implemented six tracking strategies with various starting time points and durations to capture potential intermediate differentiation events (Figure 3A). Single-cell transcriptome analysis revealed that the EBs differentiate into six main cell types: the primordial germ cell-like (PGC-like) cells and cells from all three germ layers including neuron, gut, heart, endothelium, blood and cardiopharyngeal mesoderm (CPM). We observed a mesoderm-preferred differentiation in this EB system, particularly for heart lineages (Figures 3B and Figure S3 B-F).

**Figure 3.**
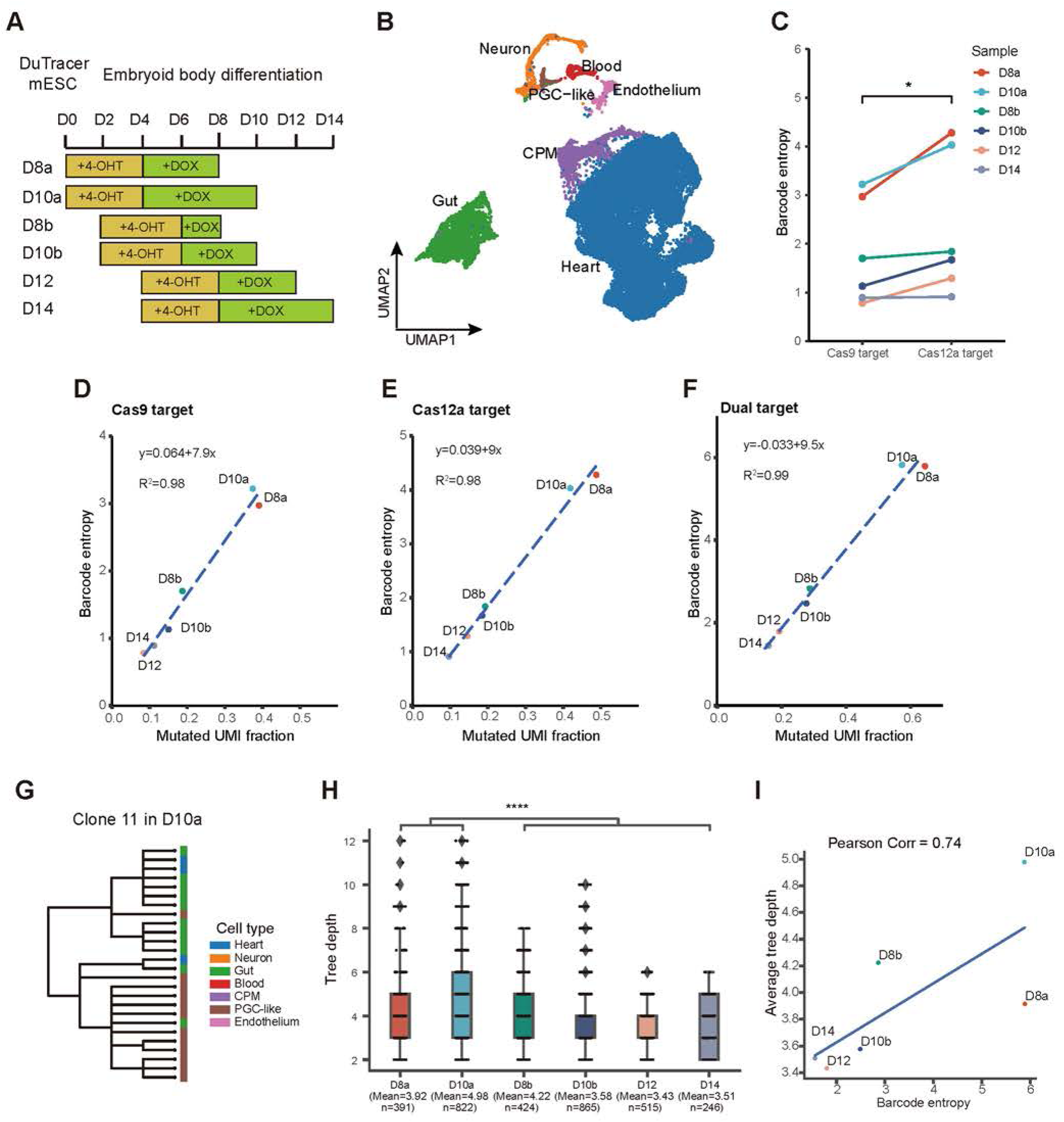
A determined barcoding system may have constrained barcoding capacity. (A) Experimental design of induction strategies in mouse embryoid bodies (EBs). (B) The cell type annotation of integrated scRNA-seq of six EB samples. PGC-like, primordial germ-like cells; CPM, cardiopharyngeal mesoderm. (C) Comparison of the barcode diversity between Cas9 and Cas12a targets (paired two-tail t-test is indicated: *p < 0.05. (D-F) The relationship between barcode entropy and mutated UMI fraction. Cas9 target (D), Cas12a target (E), dual target (F) (G) An exemplary reconstructed phylogenetic tree. The leaves are colored according to cell types. (H) The tree depth distribution of six EB samples (unpaired two-tail t-test is indicated: ****p < 0.0001). (I) The relationship between the barcode diversity and the average tree depth.

We observed minimal deleterious inter-site deletions in this EB differentiation model (Figure S4A-D), further demonstrating that DuTracer retains more recorded information and maintains higher lineage recording fidelity. Barcode entropy is a summary metric of the average barcode diversity, considering both the numbers and their frequency of indel alleles ^24^. We found that Cas12a gene editing showed higher barcode diversity than Cas9 in the current implementation as evaluated by barcode entropy (Figure 3C). This result indicates that the Cas12a gene editing system may retain unique features to serve as a promising toolkit for future development on lineage tracing, as previously reported ^24^.

With these varied recording strategies, we were able to evaluate the mutation patterns under different scenarios. We discovered a strong linear correlation between barcode entropy and the UMI (unique molecular identifier) fraction of mutated alleles for targets of Cas9, Cas12a, and both (Figure 3D-F). This linear correlation was also observed for each CRISPR target (Figure S4E-H). Using the fitted linear function, we could estimate the maximum barcode entropy once all targets are mutated (mutated UMI fraction = 1). These analyses indicate that DuTracer has a theoretical upper-bound barcoding capacity, which appears to be an intrinsic property of gene editing on targets for Cas9/Cas12a and their combinations.

Despite inevitable cell dropout and barcode dropout, we identified numerous clones in each sample based on combined intBCs (Figure S4I-L). Within each clone, a phylogenetic tree of different cells could be reconstructed to infer the lineage coupling. For example, the detailed relationship between PGC-like cells, heart, and gut could be visualized at the single-cell level for clone 11 in sample D10a (Figure 3G). Notably, we found that the average tree depths of lineage trees were deeper in D8a and D10a than in other samples (Figure 3H). Given that the barcode entropy was also higher in these two samples, we explored the potential connection between barcode entropy and the depths of reconstructed trees. Our analysis confirmed a positive correlation (Figure 3I). Therefore, DuTracer highlights an upper limited tree depths for single-cell lineage recording due to the constraint of maximum barcode entropy.

### Information gain reveals cell determination events

After evaluating the basic properties of the barcoding system, we sought to dissect the recorded cell determination events. Trajectory analysis using the CytoTRACE ^36^ tool provided insights into cell state transitions within cell types (heart and neuron) but had difficulties in revealing the developmental relationships among different cell types (Figure S5 A, B). In contrast, clonal analysis integrating barcode information and cell transcriptomes offers a unique perspective to delineate the lineage relationships among cells ^21^. To reveal the clonal relationship, we assigned the cells into clones and subclones based on the shared intBCs (static) and CRISPR indels (cumulative), respectively. At the clonal level, we observed that most of the clones contained all six cell types, albeit varied compositions of each cell type (Figure 4A and Figure S5C). This result indicates that the overall developmental potency bias toward heart lineage of the EB model did not occur at the initial stage of cell aggregation when the clone was labeled, but rather during the subsequent cell differentiation. To investigate the lineage at the subclonal level, we reconstructed the phylogenetic trees for each clone using Cassiopeia ^37^. To improve the visualization of the relationships among cells within individual samples, we connected the cell phylogenetic trees from various clones to a pseudo-root (Figure 4D-I). The integrated phylogenetic trees revealed that diverse cell specification events can be captured between samples, which is presumably attributed to the utilization of multiple gene editing strategies and the stochastic nature of gene editing.

**Figure 4.**
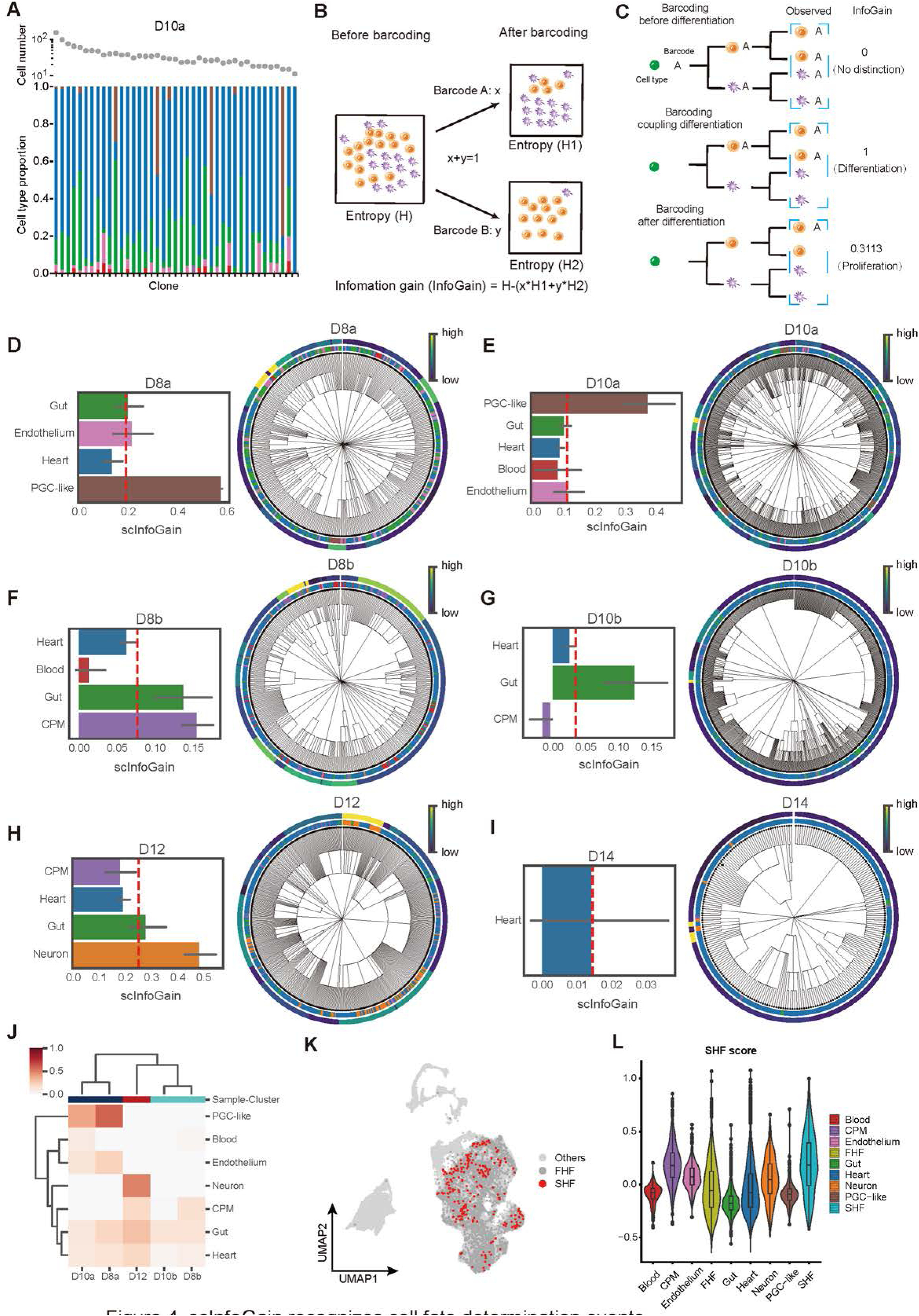
scInfoGain recognizes cell fate determination events. (A) Cell-type distribution of called clones. (B) Illustration of the information gain (InfoGain) calculation. x and y are the proportion of cells labeled by barcodes A and B, respectively. (C) Models show the relationship between the InfoGain and the barcoding process. InfoGain is calculated based on the observed barcoding cells in a dashed square. A, barcode. Different colors represent different cell types. (D-I) Distributions of scInfoGains for various cell types in six EB samples and their corresponding pseudo-phylogenetic trees. D8a (D), D10a (E), D8b (F), D10b (G), D12 (H), D14 (I). Red lines represent average scInfoGains, and error bars represent 95% confidence interval. Each pseudo-phylogenetic tree has two circles. The inner one exhibits cell types, and the outer one displays scaled scInfoGain values, as indicated by a gradient color bar. (J) Hierarchical clustering of scInfoGains for eight cell types. Sample D14 was removed for its existence of only one cell type. (K) UMAP distribution of potential FHF and SHF inferred from clonal coupling. (L) Violin plot of SHF scores for various cell types.

Given this vast diversity, accurately identifying lineage branching points within the tree topology would be challenging. To address this, we adapted the concept of information gain (InfoGain), which has been typically used to assess the coupling between features and classification results ^38–40^, to compute the coupling between barcodes and cell types (Figures 4B, C; Methods). Based on a theoretical model of barcoding processes, early barcoding indiscriminately labeled all cell types and failed to distinguish differentiation events within progenies. Accordingly, no coupling was observed between any specific cell type and barcodes, resulting in an InfoGain of 0. In contrast, precise barcoding exactly at the branching points produced the highest InfoGain, which was equivalent to the entropy of cell-type distribution. Even late barcoding during the post-branch proliferation stage enabled the recognition of fate-determination events, albeit with reduced coupling strength between barcodes and specific cell types (Figure 4C).

In essence, a strong coupling indicates that barcoding occurs soon after the divergence of cell types. We hypothesize that InfoGain may provide a rough estimation for the timing of differentiation, i.e. branching, based on the timing of barcoding inferred through the phylogenetic trees. Inspecting the D10a sample that has the highest average tree depth, we observed distinct lineage segregation of PGC-like cells in a specific clone (Figure S6A). By comparing the InfoGain of the adjacent level of barcodes within this clone, we could efficiently identify the critical lineage segregation points labeled by barcode II, suggesting that cell types diverge around the time of barcode II generation. Additionally, intra-clonal feature selection of PGC-correlated genes for cells within barcode II-labeled clones indeed revealed key PGC fate determination genes, including *Dppa5a* (Figure S6A; Methods). These findings underscore the potential utility of InfoGain as a metric for identifying critical lineage segregation points and inferring their branching time.

Based on InfoGain, we developed a refined metric named scInfoGain to effectively assess the overall barcoding ability of a phylogenetic tree in recognizing the lineage divergent events (Figure S6B; Methods). scInfoGain leverages the InfoGain values from all levels of barcodes associated with a particular single cell, providing an average quantification of the degree to which barcoding captures cell differentiation events. Evaluating the scInfoGain across different cell types in all six samples revealed intriguing patterns (Figure 4D-I). We found that PGC-like cells exhibited high scInfoGain levels in samples D8a and D10a; the gut showed relatively elevated scInfoGain in samples D8b, D10b, and D12; neurons displayed relatively high scInfoGain in samples D12 and D14; whereas heart cells consistently showed low scInfoGain levels in all samples. We noticed that in all respective samples, PGC-like cells were specified earlier than cells from mesendodermal cells derived from the epiblast, while endoderm-derived gut and ectoderm-derived neurons were specified earlier than the mesodermal subtypes. These results suggest that cell types segregated earlier tend to display higher scInfoGain levels. Additionally, we found blood cells exhibited a high level of scInfoGain in sample D8a but lower levels in sample D10a and D8b (Figure 4D, E), suggesting that the stochastic nature of gene editing may affect the characterization of the cell fate determination process. Moreover, we extended our analysis to a mouse embryo lineage-tracing dataset ^30^, where the scInfoGain metric effectively identified early-specified cell types, such as visceral endoderm at the E8.5 stage (Figure S6C, D). Together, scInfoGain quantifies the overall extent of differentiation recorded for each cell type and may be applied to pinpoint the early segregated cell types of a divergent process.

Distinguishing cardiac cells originating from the first heart field (FHF) and the second heart field (SHF) based on canonical markers proved challenging in single-cell transcriptomic analysis. However, cell type clustering based on scInfoGain revealed that CPM and gut cells had a close lineage relationship with the heart (Figure 4J). Cardiac cells are known to derive from two major lineages: the FHF, originating from early lateral plate mesoderm near the foregut, and the SHF, arising from the neural crest, which transits through CPM ^41–44^. Given the cell division history tracked by the DuTracer recorder, we hypothesized that cardiac cells that are clonally connected to CPM may originate from SHF. Accordingly, we classified cardiac cells from clones containing more than 18% CPMs as SHF-derived cells, while those from the other clones were considered FHF-derived heart cells. To avoid potential confounding influences, we did not categorize those heart cells not in clones (Figure 4K). Compared to FHF, SHF displayed dozens of differentially expressed genes (Figure S6E). By using these up-regulated genes in SHF as gene signatures, we confirmed their high expression activity in both CPM and SHF (Figure 3L), indicating our clone-based approach aids in identifying nuanced yet enduring transcriptional signatures that discriminate their distinct origins among closely related cell types.

### Information gain quantifies the overall barcoding capacity of DuTracer in differentiation models

In addition to coupling analysis focused on specific cell fate determination events, we realized that InfoGain could be adapted to quantify the degree of differentiation events distinguished by a set of non-overlapped barcodes. For instance, in an ideal labeling experiment, when the total InfoGain of barcode set A-D equals the entropy of cell-type distribution, barcode set A-D will effectively distinguish each cell type. Conversely, if the barcode sets at the same barcoding level contain only barcodes A and B, not all cell types are distinguishable, and the total InfoGain is less than the entropy of cell-type distribution (Figure 5A). This implies that when the total InfoGain is below the entropy of the cell-type distribution, additional barcodes are needed to accurately label the differentiation process.

**Figure 5.**
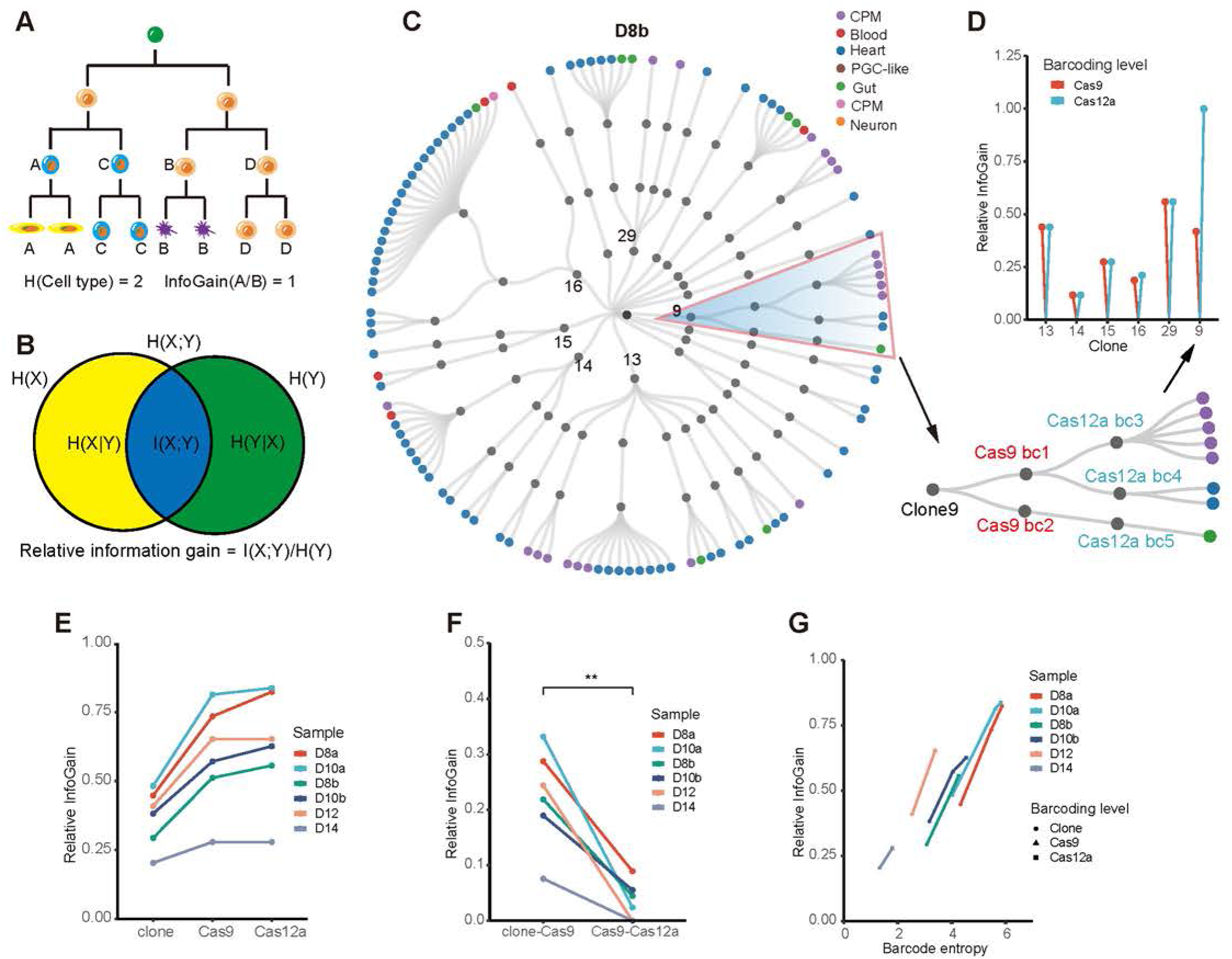
RelativeInfoGain quantifies the extent of barcoding. (A) A model shows the calculation of the information gain for a barcode set. (B) Definition of relative information gain (RelativeInfoGain). X represents a random variable denoting the distribution of barcodes, and Y represents a random variable denoting the distribution of cell types. H(X) is the entropy of the barcode variable, and H(Y) is the entropy of the cell type variable. I (X;Y) is mutual information of X and Y, which also represents the information gain of X in the context of Y. (C) A representative example shows the calculation of RelativeInfoGain in Cas9 and Cas12a levels using clone 9. Internal nodes of the circular sparse lineage plot from center to periphery represent samples, clones, Cas9 barcodes, and Cas12a barcodes, respectively. Terminal branches are cells labeled according to their cell type designation. (D) Changes of RelativeInfoGains in Cas9 and Cas12a clonal levels. (E) RelativeInfoGain across three clonal levels of the barcoding process. (F) Comparison of the increment of RelativeInfoGains between sequential barcoding processes (paired two-tail t-test is indicated: **p < 0.01). clone-Cas9, RelativeInfoGain difference between clone level and Cas9 level. Cas9-Cas12a, RelativeInfoGain difference between Cas9 level and Cas12a. (G) Line plot shows the relationship between barcode entropy and RelativeInfoGain. The slope of the line indicates the RelativeInfoGain encompassed by per unit of barcode entropy. Note: The barcode entropy of Cas9 is the combined diversity of intBC and Cas9 indels within the same alleles, while the barcode entropy of Cas12a is the combined diversity of intBC, Cas9 indels, and Cas12a indels within the same alleles. Two samples (D12 and D14) don’t display increasing barcode entropy and corresponding RelativeInfoGains from the Cas9 level to the Cas12a level.

To facilitate the comparison of the labeling process between different experiments and aid the further design of the barcoding strategy, we normalized the InfoGain against the total entropy of cell-type distribution (referred to as RelativeInfoGain) (Figure 5B). We adapted the RelativeInfoGain metric to assess the barcoding property of the DuTracer system in three adjustable barcoding levels: the clone level, labeled by intBCs at the initial stage; the Cas9 level, marked by Cas9 indels following 4-OHT induction; and the Cas12a level, barcoded by Cas12a scars following doxycycline (Dox) induction. As an example, we observed that the increase of RelativeInfoGain across the sequential level of barcodes was consistent with the finer labeling of cell types by an increasing number of corresponding barcodes in clone 9 of sample D8b, by calculating the RelativeInfoGain of a barcode set on the same barcoding level (Figure 5C, D). In addition, a comparison of RelativeInfoGains among different clones could effectively reveal the total extent of distinguished cell types and their changes across two gene editing levels (Figure 5C, D).

We further analyzed the RelativeInfoGain change among different barcoding levels in these six EB samples (Figure S7A). Using permutation simulations, we validated that the observed RelativeInfoGain captured the coupling information between barcode sets and cell types (Figure S7 B). A comparison of the RelativeInfoGain across different stages of barcoding revealed that the Cas9-early barcoding recorded more differentiation events than Cas12a-late barcoding, indicating that a considerable number of differentiation events were recorded in the early stages of EB development (Figure 5E, F). Nevertheless, we found that Cas12a-late barcoding exhibited RelativeInfoGain values comparable to the Cas9-early barcoding, with respect to per unit barcode entropy (Figure 5G). As the change of barcode entropy between consecutive barcoding processes reflects the capability of the latter stage to generate unique barcodes, this result indicates the reduced number of differentiation events recorded during the late stage was primarily attributed to the decreased ability to produce non-redundant barcodes. Collectively, our findings demonstrated that the sequential barcoding by two nucleases effectively captures crucial differentiation events throughout various developmental stages, and the extent of this barcoding ability can be quantified using RelativeInfoGain.

### DuTracer facilitates the discovery of fate-biased molecular drivers

Neuromesodermal progenitors (NMPs) are known to differentiate into the posterior spinal cord and paraxial mesoderm ^45–48^. This heterogeneous population of progenitors displays differentiation bias towards mesoderm or neural cells ^45,49,50^. However, the molecular mechanisms regulating this fate bias remain elusive. Single-cell lineage tracing technologies offer an opportunity to uncover the refined mechanisms crucial to cell fate preference. For example, by simultaneously analyzing the clonal transcriptomes at both early and late stages, researchers have demonstrated the feasibility of probing lineage-biased regulations *in vitro* or *ex vivo* ^51^.

To unveil the molecular drivers governing cell fate decisions during NMP differentiation, we implemented DuTracer recording in a mouse neuromesodermal organoid (NMO) model. Following an approach similar to that used in human NMO studies, we first derived NMPs from mouse ESCs and allowed these NMPs to self-organize and form mouse NMOs (Figure S8A, B). As most cell types emerged around D30 in human NMO, we first analyzed the lineage and transcriptome of the mouse NMO at the same time point. The scRNA-seq showed both neural cell types and mesodermal cell types were successfully generated (Figure S8C). However, after analyzing the lineage coupling of different cell types, we found that fewer than 300 cells were called in the phylogenetic tree and most clones were biased for specific cell types. Notably, neural cells were underrepresented in the lineage data compared to their transcriptomic presence (Figure S8D), likely due to the silencing of barcode expression in neural lineage after prolonged culture (Figure S8E).

To avoid possible transgene silencing and build the state-fate connections between NMPs and their derivatives, we collected the clonally related cells from both early NMP progenitors (D3) and late NMOs (D15) (Figure 6A). Single-cell transcriptome analysis showed that D3 cells were primarily NMPs, and D15 cells consisted of spinal cord neural cells and mesodermal cells (Figure 6B, C and Figure S9A). Of note, the D15 sample displayed comparable barcode expression in both neural and mesodermal cell types (Figure S9 B). Meanwhile, we observed a decline in the number of intBCs per cell at the late stage of organoid development (Figure S9C). Combined with NMO D30 observations, these results suggest that transgene expression is indeed suppressed over time.

**Figure 6.**
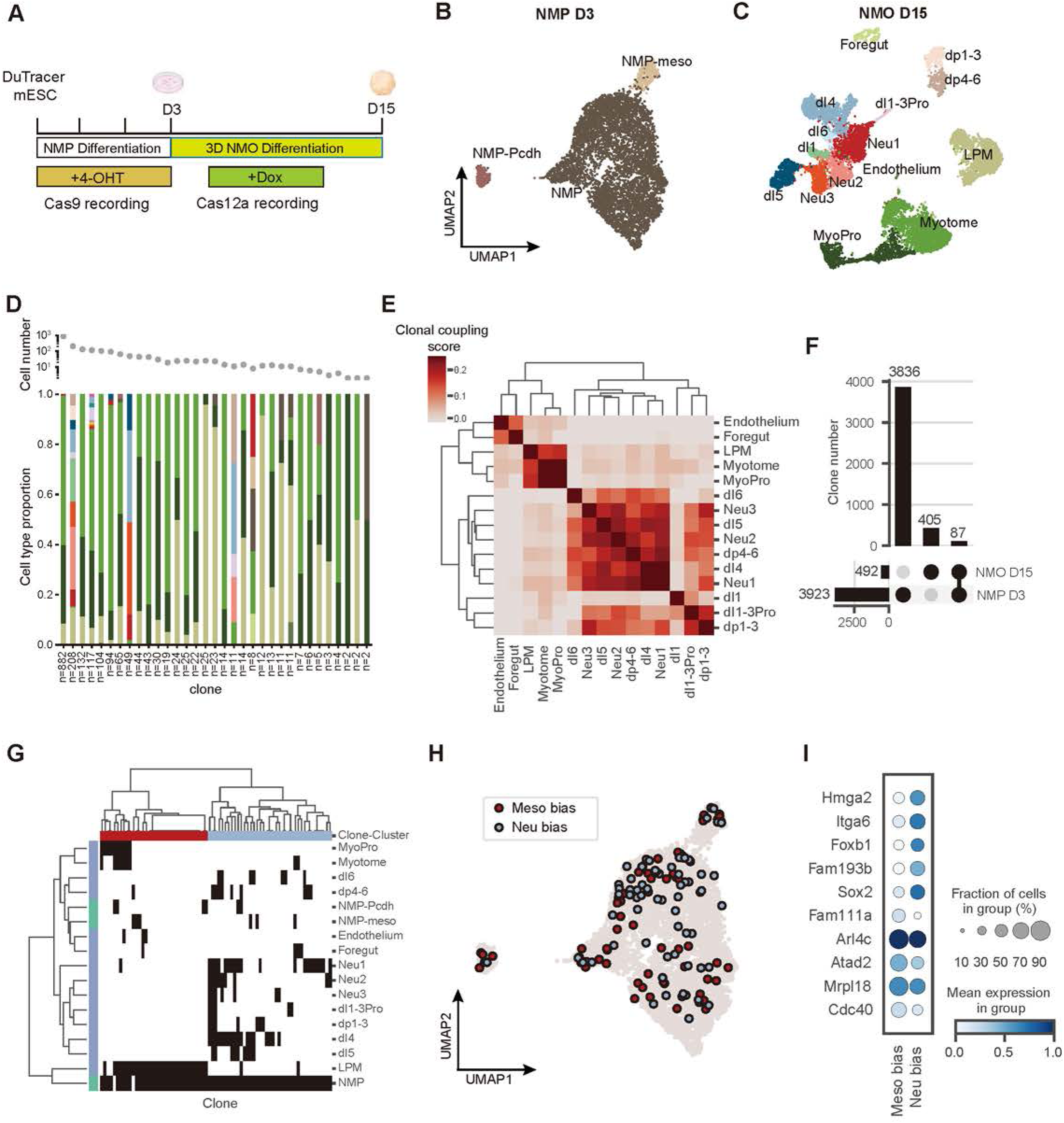
Lineage tracing reveals the fate bias of NMPs. (A) Experimental design of cross-timepoint experiments. (B-C) Major cell types in NMP D3 (B) and NMO D15 (C). NMP-meso, NMP with mesoderm features; NMP-Pcdh, NMP expressing protocadherin genes; NeurPro, neural progenitors; MyoPro, myocyte progenitors; LPM, lateral plate mesoderm; Neu1-3, unclassified neurons. (D) Cell-type distribution of called clones. Only clones containing myogenic lineages are shown. Cell type colors are shown according to Figures B and C, and cell numbers of each clone are indicated. (E) Clonal coupling among different cell types. (F) Upset plot of clones in early and late stages. (G) Cell-type distribution of 87 shared clones. (H) UMAP distribution of fate-biased NMP D3 progenitors of 87 shared clones. (I) Selected differentially expressed genes between mesoderm-biased and neuro-biased NMP D3 progenitors.

By integrating clonal information with single-cell transcriptome data, we evaluated the clonal relationship of cells within the D15 NMO, and between D3 and D15. Despite attempts to delineate the lineage segregation points between different cell types within D15, a lack of mutation diversity hindered the reconstruction of an informative phylogenetic tree (Figure S9D) albeit rare inter-site deletions (Figure S9E), suggesting that mutations primarily occurred at the early stages of organoid development, before extensive cell progenitor differentiation.

We treated CRISPR scars as static barcodes and defined unique combinations of intBC and CRISPR indel as clones, aiming for a more accurate depiction of the clonal relationship between cells across the D3 and D15 stages. We found that the average clone size at D15 was larger than that at D3, indicating a proliferative cell division (Figure S9F, G). Remarkably, we observed that the cell fate bias in NMOs was apparent at the clonal level in D15. An exemplary clone contained nearly 900 cells, most of which were myotome and myoblasts (Figure 6D). This myogenic lineage bias was also evident in one clone from the D30 sample (Figure S8F). Additionally, we computed the lineage coupling scores of different cell types (Methods). Clustering analysis based on the scores identified two major clusters: one enriched in neural cells and the other predominated by mesodermal (Figure 6E). Therefore, DuTracer effectively captures the major cell fate divergences of neuromesodermal lineages.

Next, we visualized the cell relationship within individual clones across both the D3 and D15 stages. Approximately 4,000 clones were identified at the D3 NMP, while around 400 clones were found at the D15 NMO (Figure 6F). Only 87 clones were shared between the early and late stages of the organoid differentiation. Additionally, uneven expansions were observed in the shared clones (Figure S9H), implying that clone dominance occurred along NMO development. Upon grouping these shared clones, we observed distinct compositions of cell types, which were further categorized into two major neural and mesodermal clades (Figure 6G). By mapping the D15 cell fates to their possible progenitors at D3 for individual shared clones, we analyzed the state difference between neural-biased and mesoderm-biased D3 cells (Figure 6H). Our analysis identified a number of genes differentially expressed between the neuro-biased and mesoderm-biased progenitors (Figure 6I). Specifically, *Sox2* and a few other genes, including *Foxb1*, exhibited higher expression levels in neuro-biased cells. *Sox2* is a well-known driver of neural lineage development, and *Foxb1* has also been found essential for posterior spinal cord development in Xenopus ^52^. These results support the DuTracer’s ability to track clones over time, which can provide insights into progenitor state differences that influence later cell fate diversification.

## DISCUSSION

Under the current implementations of CRISPR-based lineage tracking technologies, tandem targets within a short DNA fragment are subjected to rapid target exhaustion and potential inter-site deletions spanning multiple targets. To circumvent these limitations, DuTracer exploits two gene editing systems to separate the nuclease-cutting events into two independent stages. This strategy greatly reduces the rate of target dropouts and enhances the recording intervals, enabling the labeling of more cell divisions, and consequently, more differentiation events as quantified by information gain. We noticed that an application of orthogonal nucleases in lineage tracing has been demonstrated in the zebrafish model. However, this method still risks large inter-site deletions due to its arrangement of multiple target sites within a locus ^53^. We confirmed the veracity of low inter-site deletion of DuTracer in various cellular models and highlighted the benefits of a deeper lineage tree, and higher resolution conveyed by this system.

In addition to increasing the fidelity of the barcoding process, we realized that DuTracer could serve as a tangible system for comparing gene editing properties and interactions between these two most popular CRISPR nucleases within the same cells. In these two barcoding systems and their combinations, we demonstrated that increasing the number of targets by inserting additional copies of the target array may enhance barcode diversity for recording cell divisions. However, the diversity would reach saturation once the copy number surpassed certain thresholds. Furthermore, the saturation speed appeared to be affected by mutation rates. Interestingly, we discovered that the barcode entropy highly correlates with the mutated UMI fraction, and it appears that the barcode entropy might be maximized when the mutated UMI fraction reaches 100%. These results imply that once a barcoding system is established, it inherently possesses a maximum barcoding capacity. Moreover, our findings indicated that the average reconstructed tree depths are associated with barcode entropy. As barcode entropy may be fixed for a given recorder, the average reconstructed tree depths would also have an upper limit. Thus, the lineage recorder should be carefully chosen to provide sufficient barcode entropy for the process under study. In our EB models, we demonstrated that adjusting the release of barcoding entropy influences the characterization of different events. To maximize a recorder’s utility in a specific differentiation model, gene editing should be controlled to record as many differentiation events as possible. Our work also underscores certain guidelines for designing lineage recorders, such as using lineage recorders with CRISPR targets exhibiting diverse indels and adequate mutation rates, as also indicated by previous studies ^24,30,33^.

We provided information gain to precisely quantify the coupling relationship between the barcoding process and cell differentiation. This metric can be used to quantify the recording efficiency of a barcoding system or to reveal the extent of lineage specification labeled by the barcoding system. The former objective, evaluated by RelativeInfoGain, provides directions for refining barcoding strategies, such as adjusting the timing of gene editing induction. For example, we could induce gene editing of both nucleases at the early stage of EB development to capture more differentiation events. The latter goal, quantified by scInfoGain, leverages information gain across all subclonal layers. Through calculating scInfoGain, we can gain insights into the differentiation patterns of a specific cell type, and further infer the lineage specification order and lineage relationships among different cell types. We demonstrated that this metric was generally applicable to other CRISPR/Cas9-based lineage tracing system used in mouse embryogenesis ^30^. However, it is important to note that both RelativeInfoGain and scInfoGain calculations are influenced by the complex interplay of biased cell differentiation and stochastic gene editing. High barcode diversity and diverse cell type generation are favorable when applying scInfoGain-based analysis.

Single-cell lineage tracing technology holds the promise of elucidating the lineage history of specific cell groups, the hierarchy of cell potency, and potential molecular drivers of cell fate determination. Our scInfoGain-based analysis indicates that the differentiation order of cell types in mouse EBs may also follow the same principle as their *in vivo* counterparts. Specifically, we demonstrated that clonally related CPM and SHF shared heritable transcriptional features. Beyond randomly differentiated EB models, we have also applied DuTracer to a more constrained differentiation system—mouse neuromesodermal organoids. By clonal tracking, we showed that individual NMP clones exhibited strong cell fate bias. Furthermore, mapping the state-fate relationship of cells in early and late stages identified a set of candidate genes potentially involved in regulating cell fate choices.

Finally, considering the feasibility of organoid generation and barcode introduction, further application of DuTracer in human organoids may facilitate the discovery of the unique features of human development. We also anticipate that further optimization of DuTracer guided by detailed evaluations of barcode entropy and information gain will make this technology a powerful tool to record cell lineages in development or tumorigenesis.

### Limitations of the study

The low rate of gene editing for some EB samples may arise from the unexpected transgene-silencing of nucleases, a common issue of random integration ^54^. Additionally, some analyses such as hierarchical clustering of cell types and information gain calculation were based on limited sampling of cells, which may miss some coupling information of cell types and cause potential errors. Furthermore, the current application of DuTracer suffered greatly from editing leakages, thus violating the assumption that nucleases only record differentiation events after induction. Given the known leakage of induction systems (Tet-on and 4-OHT-ERT) ^34,55^, it is imperative to intensify endeavors aimed at diminishing the leakage rate to facilitate effective sequential gene editing. Finally, our DuTracer method is currently utilized in *in vitro* systems, the performance of this method remains unclear in the *in vivo* mouse model. A DuTracer animal model is eagerly anticipated.

## ACKNOWLEDGMENTS

We are grateful to Dr. Liangxue Lai for sharing the Cas12a crRNA plasmid, Dr. Yu Wang for sharing the HIT-Cas9 plasmid, and Dr. Jonathan Weissman for providing the Cas9 gRNA sequences. This work was supported in part by National Key R&D Program of China (2018YFA0801402), National Natural Science Foundation of China (32270854, 32161160322), Guangdong Basic and Applied Basic Research Foundation (2019B151502054), Science and Technology Planning Project of Guangdong Province (2023B1212060050, 2023B1212120009), Basic Research Project of Guangzhou Institutes of Biomedicine and Health, Chinese Academy of Sciences (GIBHBRP23-01), Major Project of Guangzhou National Laboratory, Grant No. GZNL2023A03005. C.C. was partly supported by National Natural Science Foundation of China (U21A20203 and 32300679), Guangdong Basic and Applied Basic Research Foundation (2022A1515110942), and Science and Technology Program of Guangzhou (2024A04J4274).

## AUTHOR CONTRIBUTIONS

G.D.P. conceived and supervised the entire study. C.C. proposed the initial design of DuTracer. Y.X.L. and C.C. designed the experiments. Y.X.L. conducted the experiments. C.C. and Z.M. performed the bioinformatic analysis. L.W., X.R.Y., M.S.L., contributed to the experimental data collection. C.C., G.D.P., Y.X.L, and Z.M., wrote the manuscript.

## DECLARATION OF INTERESTS

The authors declare no competing interests.

### Materials availability

Plasmids and cell lines generated in this study about DuTracer are available from the lead contact upon request.

### Data availability

The accession numbers of this paper are NCBI GEO: GSE261211 and GSE236936.

### Code availability

The instructions and codes to generate figures in this paper can be found at (https://github.com/gpenglab/DuTracer)

## METHODS

### Cell Lines

HEK293T cells were cultured in DMEM basic media supplemented with 10% FBS. mESCs were cultured on plates coated with 0.1% gelatin in DMEM high-glucose media containing 15% FBS, 1 mM sodium pyruvate, 1 mM non-essential amino acids (NEAA), 1% GlutaMAX, 0.1 mM 2-mercaptoethanol, 1000 U/ml leukemia inhibitory factor (LIF), 3 μM CHIR99021, and 1 μM PD0325901. All the cell lines were cultured in humidified incubators at 37*℃* in 5% (v/v) CO_2_.

### DuTracer plasmid design and cloning

The DuTracer system includes three components: (1) Dual-nuclease plasmid: A PiggyBac plasmid comprising a Cas9 coding region fused with two ERT2 domains, driven by the EF1α promoter, and a Cas12a coding region controlled by Tet-ON promoter. The Cas12a fragment was cloned from the pY016 plasmid (Addgene: #69988) and the Cas9-ERT2 part was cloned from the HIT-Cas9 plasmid (a gift from Dr. Yu Wang). (2) Target plasmid: A lentiviral plasmid with constitutive mCherry expression driven by EF1α promoter. A fragment containing four targets (synthesized by IGE) was inserted in the 3’UTR after the mCherry coding region. The target plasmid was derived from the pSin-EF1α-Flag vector (a gift from Dr. Hongjie Yao) with the following modification: The flag sequence was replaced by the mCherry-Target sequence using Gibson cloning, and the UCOE (ubiquitous chromatic opening element) sequence (synthesized by Sangon Biotech) was inserted before the EF1α promoter to reduce the gene silencing. NheI and BamHI sites were placed before the target region to facilitate the addition of integration barcodes. IntBCs were inserted into the plasmids using the NEBuilder HiFi Assembly Master Mix (NEB). (3) gRNA plasmid: A lentiviral plasmid used for constitutively expressing two Cas9 sgRNAs and two Cas12a crRNAs (a gift from Dr. Liangxue Lai) driven by separate human U6 promoters. The synthesized gRNA sequences were added to the Psin-EF1α-BFP plasmid using Gibson cloning.

### Lentiviral packaging

All lentiviruses were produced in HEK293T cells using the second-generation lentiviral system. Before plasmid transfection, 1× 10^7^ HEK293T cells were plated in a 100 mm dish. Plasmid transfection was performed when the cells reached 70-90% confluency. For transfection, 6 μg Target plasmids, 4 μg PSPAX2, and 2 μg PMD.2G plasmids were used plus with 48 μl PEI. After 12 hours, the transfection reagent was removed by washing the plate with a fresh medium. After 48 hours, the lentiviral suspension was collected and filtered through a 0.45 μm filter to remove cell debris.

### Generation of DuTracer cell lines

We used the PiggyBac transposon system and lentiviruses to construct DuTracer cell lines (HEK293T or mESCs). Cells were transfected with 2 μg of dual-nuclease plasmid and 0.66 μg of transposase plasmid using Lipofectamine 3000 reagent (Thermo Fisher) and positive clones were selected based on their resistance to puromycin. Subsequently, survived cells were infected with gRNA lentiviruses. Three days post-infection, fluorescence-activated cell sorting (FACS) was utilized to isolate BFP-positive cells. Finally, the enriched BFP-positive cells were further infected with a high MOI Target lentivirus to achieve a high copy number of insertions, and mCherry-positive cells were isolated using FACS.

### Bulk amplicon sequencing

For mouse ESCs, the DuTracer cell line was established and seeded at an appropriate density, denoted as Day 0 (D0). Every two days, the cells were dissociated into single cells, with half of the cells being used for DNA extraction and the remaining half reseeded for subsequent induction. Throughout the D0-D8 culture period, 200 nM 4-OHT was continuously supplemented to induce Cas9 activity. From D8 to D16, 2 μg/mL Dox was added to induce Cas12a activity. ESC samples were collected at D2, D4, D6, D8, D10, D12, D14, and D16. Similarly, HEK293T samples were collected at D1, D3, D5, D7, D8, D10, D12, and D14, with dox induction for seven days, followed by 4-OHT induction for another seven days.

Genomic DNA was extracted using the HiPure Tissue DNA Micro Kit (Magen). The barcode region was amplified via nested PCR. A first-round PCR was conducted with 500 ng of gDNA per reaction, and four parallel PCR reactions were performed for each sample to minimize PCR bias. The PCR product was combined and purified using 0.7x SPRI beads. Approximately 100 ng of the first-round PCR product was used in a second-round PCR to introduce Illumina adaptors and indexes. The PCR products from the second round were purified using 0.8X SPRI beads, followed by sequencing on the Illumina NovaSeq6000 platform with PE250 settings.

### Mouse embryoid body differentiation

We used the traditional hanging drop method to generate embryoid bodies as previously described ^35^. Briefly, mESCs were dissociated into single cells by incubating with 0.25% Trypsin for 5 min, then the cells were suspended in mESC media without 2i and LIF at a cell density of 3×10^4^/ml. After adding 10 ml PBS to the bottom of the dish, we used the multichannel pipette to add 30 μl drop on the lid of the dish and then carefully placed the dish in the incubator. After EBs formed and grew for three days, they were transferred to 6-well plates coated with low-attachment reagents (STEMCELL). The differentiation medium was changed every two days until D14.

### Generation of mouse neuro-mesodermal organoids

Methods for generating neuro-mesodermal progenitors (NMPs) and organoids from mESC were adapted from previous studies ^45,56^. In brief, mESCs were dissociated into individual cells by 0.05% Trypsin, followed by plating on 0.1% gelatin precoated dish at a density of 5×10^3^ cells/cm^2^ in N2B27 medium supplemented with 10 ng/mL FGF2. After two days, the medium was replaced with N2B27 medium containing 3 μM CHIR and 10 ng/mL FGF2 for an additional day to induce NMPs.

To generate mouse neuro-mesodermal organoids (NMOs), NMPs were dissociated using Accutase to obtain a single-cell suspension. On NMO Day 0, 100 μL NMP cell suspension was plated on an ultra-low binding 96-well plate in NMO medium (N2B27 medium with 10 ng/mL FGF2, 3 μM CHIR, 2 ng/mL IGF1, 2 ng/mL HGF) at a density of 1×10^5^ cells/ml. After two days of culture, half of the medium was removed and replaced with 100 μL of NMO medium. On NMO Day 4, the organoids were cultured in N2B27 medium, with the medium being refreshed every two days.

### Single-cell RNA-seq

For single-cell preparation, EBs/NMOs were dissociated by the Papain dissociation kit (Worthington). In brief, EBs/NMOs were incubated in 1 ml Enzyme A along with 20 μl DnaseI at 37*℃* for 20-25 min, followed by gentle blowing every 5 minutes with wide-bore tips. After washing and filtering, a suitable volume of PBS with 0.04% BSA was added to resuspend the cells to achieve a cell concentration of 2×10^6^ /ml. Single-cell RNA libraries were prepared using DNBelab C4 kit (BGI-MGI) according to the user’s guide. The main steps of library preparation include droplet formation, demulsification, reverse transcription, second-strand synthesis, cDNA, and oligo product amplification following size selection. Each obtained cDNA pool was divided into two parts. One part of cDNA was prepared by fragmentation, end repair, adaptor ligation, and PCR to get a cDNA library for scRNA-seq. Another part of cDNA was used for amplicon sequencing. Corresponding oligo libraries were generated by PCR amplification from oligo products. The cDNA libraries and oligo libraries were sequenced on the MGI-seq 2000 platform.

### Single-cell amplicon sequencing

We performed amplicon sequencing to capture the DuTracer target region to obtain both the intBC and CRISPR indel information. For the single-cell cDNA template, a pair of primers (C4-F1:5’-ACACTCTTTCCCTACACGACGCTCTTCCGATCTCGTAGCCATGTCGTTCTGCG-3’, C4-R1:5’-TCTCGTGGGCTCG GAGATGTGTATAAGAGACAGTGCAGGAGCGGATTGCTTCGAACC-3’) suitable for the MGI C4 cDNA library adaptors were used for amplification. 50-100 ng cDNA templates were used per amplicon reaction, and four parallel reactions were performed for one sample to reduce potential PCR biases. PCR was performed using 2x Kapa HIFI Mix according the following program: 95*℃*, 3 min; 98*℃*, 15 s; 60*℃*, 18 s; 72*℃*, 15s. The first round of PCR was 8-12 cycles and the second round of PCR was 5-6 cycles. PCR products were selected and purified by AMPure XP beads (Beckman). The amplicon libraries were sequenced on the Illumina Nova PE250 platform or MGI-PE300 platform.

### Single-cell data analysis

Gene expression matrices were obtained using DNBelab C4 software (v1.0) with default parameters. For subsequent analysis, Seurat (v4.1.1) was employed. Briefly, cells with more than 60,000 reads and mitochondrial transcript percentage exceeding 10% were removed. Some samples displayed a bimodal distribution of mitochondrial transcript percentage, presumably originating from the sequencing of stripped nuclei. Therefore, a sample-specific minimum cutoff of the mitochondrial gene expression was applied to remove those suspected cells with stripped nuclei. Following library normalization, the top 2000 highly variable genes were identified using the variance stabilizing transformation method. The gene expression matrices were then scaled and regressed out the number of features and counts, cell-cycle score (G2M.score, S.Score), and mitochondrial gene expression. DoubletFinder (v2.0.3) was used to filter out suspected doublets post-clustering, which was conducted with a resolution of 0.8 using 30 top principal components. Subsequently, all samples were integrated, and the scaling step was repeated with the same regressing parameters using the Seurat CCA integration method. Further steps, such as principal component analysis and cluster identification, were accomplished by adjusting relevant parameters. Uniform manifold approximation and projection (UMAP) dimensionality reduction was used to project cell clusters in two dimensions. Cell annotation was first performed using a mouse reference dataset at the gastrulation stage and then carefully refined using manually curated marker genes ^42,43,57–60^.

### Calling independent clones for EB samples

We employed the modified Cassiopeia (v2.0.0) preprocessing pipeline for analyzing our amplicon sequencing data. Briefly, raw fastq files were converted to unmapped bam, low-quality reads were filtered, and bead barcodes were corrected to the provided whitelist. After collapsing the UMI sequences which differ by at most 3 bp mismatches and 2 bp indels, bead barcodes were converted to cell barcodes for further cell-level analysis. The sequences were then aligned to the target site reference, and intBCs and indel alleles were extracted from each alignment. Additionally, Starcode was utilized to correct the PCR sequencing errors by collapsing intBC within 2 bp Levenshtein distances followed by allele calling. Furthermore, UMIs within each cellBC-intBC pair were corrected to avoid allele conflicts. Clone calling was further performed after a series of filtering steps at the UMI and cell levels. Specifically, UMIs were retained if they contained at least max (5, 99th percentile/10) reads, which served as a filter for PCR sequencing errors.

### Phylogenetic tree, lineage plot, and trajectory analysis

We utilized the Cassiopeia-Greedy method to reconstruct phylogenetic trees for each clone. Obtained lineage information was then integrated with single-cell transcriptome to extract shared cells. To visualize the phylogenetic tree for all clones within a given sample, we introduced a pseudo-node to link the roots of the phylogenetic tree of each clone. The resulting single tree of shared cells for each sample was then visualized using the matplotlib library.

To integrate with the real-time information (barcoding stages according to the timing of induction), we also generated a three-layer lineage plot to display the phylogenetic relationship of cells ^61^. Lineage plots are used in RelativeInfoGain analysis for their clear visualization, easy calculation, and real barcoding intervals. We generated two types of lineage plots, dense lineage plots, and sparse lineage plots. The dense lineage plots display both the mutated and non-mutated cells, while the sparse plots only contain mutated cells with at least one indel in Cas9 targets and Cas12a targets, respectively. Unless stated, we only calculated RelativeInfoGain using the sparse lineage plots.

Trajectory analysis was done using CytoTRACE with default parameters on the integrated scRNA-seq dataset ^36^.

### HEK293T tree depth Simulation

To select the same number of targets for tree depth comparison, a maximal even number of target copies (2 N) were randomly sampled to select all the Cas9 targets (4 N) or all the Cas12a targets (4N), whereas only half of the target copies (N) were randomly sampled to extract both the Cas9 and Cas12a targets (4 N). This simulation was repeated 100 times to obtain statistical results. For simulated RelativeInfoGain of a sparse lineage plot, simulation was performed 1000 times by randomizing the barcoding relationship between barcodes and cell types for each sample.

To evaluate the effect of target insertion copies, we used the following sampling strategy to select the copy number (N) of targets: For those cells with less than or equal to N intBCs, all intBCs were selected; For those cells with more than N intBCs, only N copy of targets were randomly selected. The simulation was repeated 100 times.

### Entropy of cell-type distribution

The entropy of cell-type distribution represents the extent of cell type diversity. Cell type entropy (H) was calculated as follows:

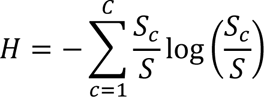

Where *S* is the total number of cells within a node, *C* is the number of cell types, *S_c_* is the number of cells of cell type *c*.

### Calculation of single-cell information gain

A similar strategy described by Yang et al. was adapted to assign InfoGain to individual cells by leveraging InfoGain from all layers of nodes ^31^. We first calculated a deviation score for each internal node in a phylogenetic tree. This deviation score was defined as the difference between the observed InfoGain and the average InfoGain resulting from random shuffling. As one internal node has one or more leaves in our trees, we selected the Laplace-corrected information gain of multiway splits, rather than the uncorrected information gain described in the main text ^62^. Random shuffle InfoGain was calculated similarly by randomly shuffling the cell type labels of each leaf under a specific internal node. This shuffling procedure was repeated 1,000 times, and the average shuffled InfoGain was used for deviation calculation. To generate the single-cell InfoGain (scInfoGain), we averaged deviation scores of internal nodes along the path from the root to a leaf and assigned the average scores to the corresponding leaf (each leaf is an observed cell). In all analyses, we filtered out cell types whose cell numbers were less than 1% of the total number of cells. Mathematically, the scInfoGain was calculated as follows:

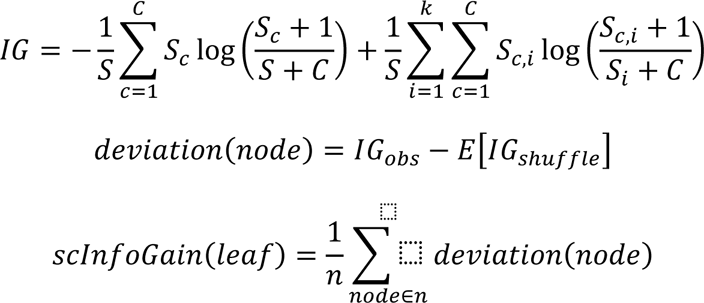

Where *S* is the total number of cells within a node, *C* is the number of cell types, *k* is the number of branch splits, *S_c,i_* is the number of cells of cell type *c* in the *k*th branch split, the “*+1*” and “*+C*” are the Laplace corrections to adjust the values affected by multiple single leaf splits.

In order to characterize the coupling relationship among cell types, the average scInfoGains for cell type were calculated within individual samples, then Ward’s hierarchical clustering was applied to the combined cellType-sample matrix.

To evaluate the correlation relationship between the branch splits and the gene expression, we utilized the “mutual_info_classif” function from the scikit-learn package, which implements a nonparametric method for detecting the correlation relationship between discrete and continuous data sets ^63^. Genes were ranked by their mutual information value and the top-ranked genes were considered candidate genes related to the branch split.

### NMO clone calling using intBC and indel combinations

To integrate static barcode and indel information to define clones in our NMO amplicon data, we first identified pre-clones using static barcodes and then refined the pre-clones by incorporating indel information to determine the final clones. Briefly, the raw amplicon data was processed as described above, and the resulting molecular table of cellBC-UMI groups was used for clone calling. The molecular table was transformed into a cellBC-by-intBC matrix *M_c,i_*, with each entry representing the UMI counts. The cellBC-by-intBC matrix was binarized at a threshold of more than 2 UMI counts. Based on the cellBC-by-intBC matrix, the cell-by-cell Jaccard similarity matrix was calculated and binarized at 0.6. This binarized cell-by-cell matrix was then converted to a graph and various graph components were identified using the Girvan–Newman algorithm. After identifying the unique intBC set for each graph component, we generated a binary intBC-by-component matrix *K_i,p_*, where each entry indicates whether an intBC is present in a specific component. To recover cells with partial dropout of static barcodes, we computed an assignment matrix as *A(c, p) = M(c, i) × K(i, p)*, where the value indicated the total UMI counts of intBCs shared between a cell and a component. Cells were assigned to the component with the highest value, and all cells within the component were defined as a pre-clone. To incorporate the indel information for refining pre-clones, we computed weighted Hamming distances between cells within each pre-clone according to their mutation types within each target site. This distance matrix was then transformed into a graph to identify graph components. If multiple graph components existed in a pre-clone, the pre-clone was further divided into smaller clones according to these components. The refined pre-clones were used as the final clones for the downstream clone analysis.

### Clonal coupling analysis

We started by forming a binary cell-by-clone matrix where each entry indicates whether a cell is present in the clone. This matrix was summed across cell types for each clone to obtain a celltype-by-clone matrix. The celltype-by-clone matrix was initially row-normalized by the number of cells in each cell type and then column-normalized. After binarized at a threshold of zero, the clone coupling matrix was generated by calculating the Jaccard similarity between cell types based on the normalized matrix. Cell-type clustering was performed using Ward’s hierarchical clustering algorithm.

## QUANTIFICATION AND STATISTICAL ANALYSIS

Unpaired two-tail t-tests and paired two-tail t-tests were used for the comparison of experimental results, and the statistical significance of results was indicated in the figure legends. Pearson correlation was applied to evaluate the correlation between two continuous variables. All statistical analysis was performed with R (version ≥ 4.1) or Python (version ≥ 3.7).

